# The dramatic impact of the PER-DBT interaction on circadian timekeeping and temperature compensation

**DOI:** 10.1101/2025.02.01.636007

**Authors:** E. Nicholas Petersen, Evelyn Keefer, Hannes Ludewig, Camille Sullivan, Dorothee Kern, Michael Rosbash

**Author notes:** Correspondence —.

## Abstract

Circadian timekeeping is orchestrated by multiple clock proteins, but mechanistic understanding is limited. AlphaFold predicts novel interactions between two helical domains of the *Drosophila* PERIOD (PER) protein and its major kinase Doubletime (DBT). These PER domains, SYQ and LT, are conserved with mammals and are predicted to interact identically with the DBT ortholog casein kinase 1 (CK1). SYQ and LT mutations designed to disrupt this interface result in severe period and temperature-sensitive fly phenotypes, and two LT mutations cause unprecedented free-running periods ≥ 44 hours and temperature-sensitive changes of ∼20 hours. Further characterization of a LT mutant strain shows a severe reduction in both PER phosphorylation and PER degradation. A human PER equivalent of the same LT mutation is a much less efficient in vitro substrate for CK1-mediated phosphorylation. These observations indicate that this conserved PER-DBT interaction is critical for PER phosphorylation, circadian timing and its enigmatic temperature compensation phenomena.

## Introduction

Circadian rhythms are an evolutionarily conserved process that allows an organism to appropriately adapt its cellular and physiological functions to other cyclical patterns in the external world. These rhythms have three basic features: 1) they have a free-running period of close to but not exactly 24 hours, i.e., ∼ 24 hours; 2) they can entrain to cycling external cues such as light, which is how they become precisely 24 hours in nature; 3) the period length changes only marginally over physiological temperature ranges^1–3^. This third feature is often described as temperature-compensation, reflecting the historical hypothesis that one or more enigmatic processes adjust a temperature-modified period back to be ∼ 24 h. However, we prefer the terms temperature-insensitivity or temperature-independence, which contain fewer mechanistic implications than temperature-compensation. Central to temperature regulation and to animal circadian timekeeping more generally is a small number of conserved eukaryotic proteins, “core clock proteins,” which engage in a well-characterized transcription-translation feedback loop (TTFL, Supp. Fig. 1-1)^4–7^.

Although insight into timekeeping and temperature regulation remains limited, *Drosophila melanogaster* has featured prominently in the few relevant mechanistic studies. This is because of its genetic facility as well as its conserved circadian system and genes. For example, and unlike in mammals, there is only one fly gene encoding each core clock protein. This facilitates assaying mutant phenotypes, which often rely on protein-coding mutations within these core clock genes and inform how their encoded proteins affect molecular events. In this context, many of the canonical rate and temperature-sensitive circadian mutations are in a conserved negative regulator of the *Drosophila* TTFL, Period (PER)^8–12^ and its modifying kinase Doubletime (DBT)^8,12,13^. Hyperphosphorylation of PER occurs over many hours eventually resulting in termination of the repression phase, phosphorylation of a PER degron region and PER degradation, and eventually a restarting of the TTFL transcription phase. The process in mammals is very similar with CK1 phosphorylating PER2^14–17^. CK1/DBT contributes to the overall rate of the circadian clock in part through this series of programmed PER phosphorylation events^17–23^. They include the phosphorylation of two different PER regions (a degron and the per^SHORT^ domain in flies vs two degrons and the FASP region in mammals), the balance of which has been proposed to contribute to the temperature independence of the central clock^15,17^.

Potentially relevant to CK1/DBT-mediated PER phosphorylation and its relationship to circadian timekeeping and even temperature regulation is an understudied interaction between PER and CK1/DBT. PER has small DBT-binding domain (DBTBD), which is conserved between flies and mammals and appears to anchor the kinase to its substrate; deletion of this domain results in arrhythmicity ^24,25^. Although this indicates that circadian timekeeping requires the PER-DBT interaction, it has made surprisingly little contribution to mechanistic understanding and temperature-independence since its discovery in 2007. This is due in large part to the lack of structural information along with the large size of PER, most of which appears unstructured (Fig. 1A).

**Figure 1.**
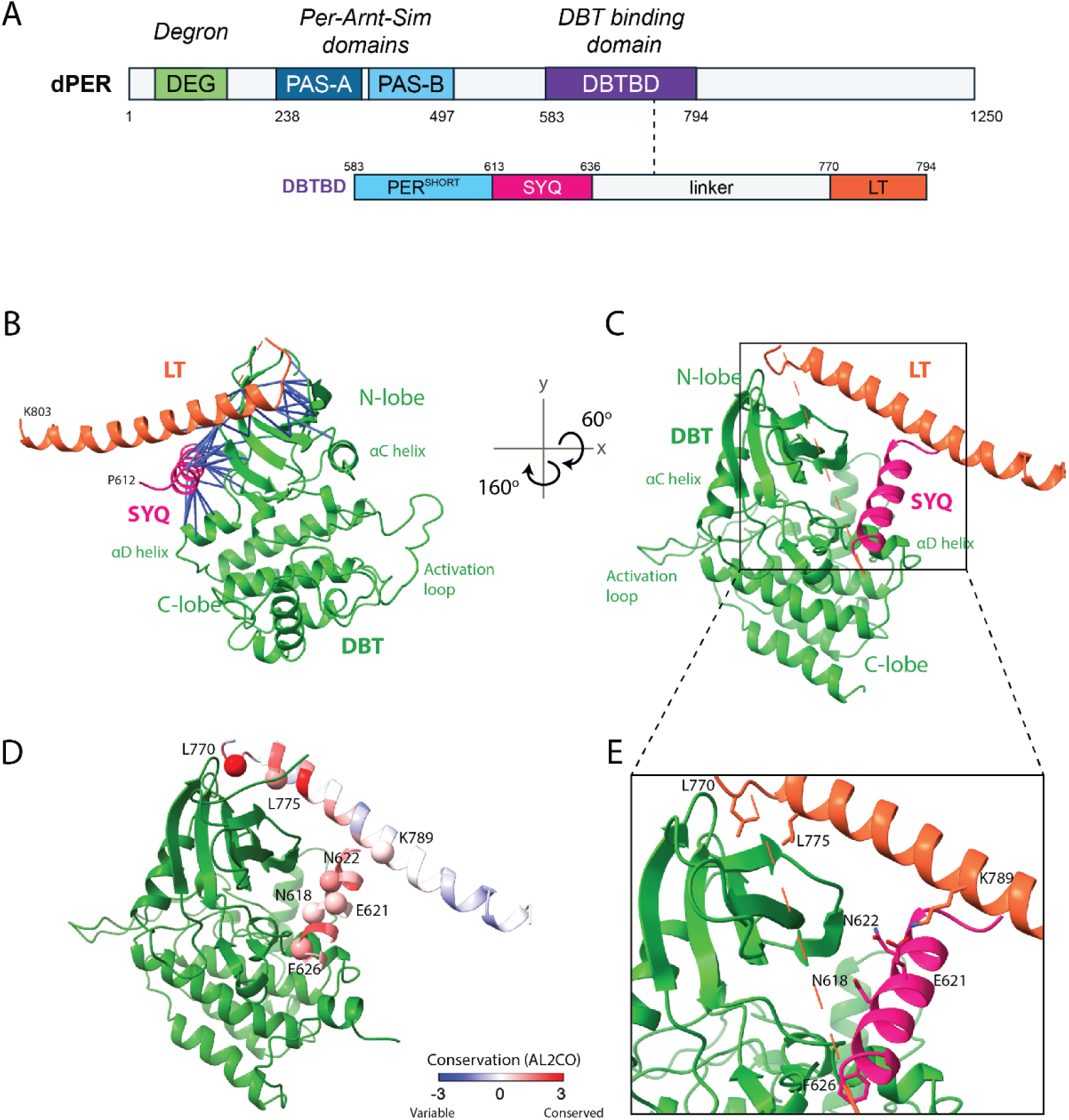
AlphaFold predicts an extensive interaction between the kinase DBT and Period LT and SYQ domains. (A) A 2D graphic showing the major domains of the PER proteins. (B) The AlphaFold prediction of the kinase DBT with the C-terminal tail hidden, with the PER LT and SYQ domains. PAE scores between PER and DBT are mapped onto the structure with a distance of 4 Å and show a high degree of confidence between these domains (blue). (C) Rotation of the complex shows that LT and SYQ sit orthogonal to one another around the N-lobe of the kinase. The location of predicted binding residues are shown in (E). (D) Conservation of the SYQ and LT domains of PER are highest (red) at the predicted interface with DBT.

Here we used AlphaFold to predict interaction details between the PER DBTBD and DBT. Mutating conserved PER amino acids that are predicted to stabilize the PER/DBT interaction results in flies with severely altered and temperature-dependent periods, demonstrating that this interaction is important for timekeeping as well as its temperature-independence. One mutation with an extremely long and temperature-sensitive period was further characterized and showed extremely slow phosphorylation, in vitro with CK1 and in vivo with DBT, as well as slow in vivo degradation along with exceptional high levels of nuclear PER in clock neurons. This not only indicates the importance of the PER-DBT/CK1 interaction to timing and temperature control but also suggests that future detailed in vitro and biophysical studies will reveal mechanistic details that will illuminate the principles that underlie circadian timing and temperature independence.

## Results

### AlphaFold Predictions

To identify regions of PER important for the PER-DBT interface, we used AlphaFold (2.3) with full-length sequences for both proteins. PER contains two well-known and structured PAS domains with neighboring helices as well as two additional predicted helices, both of which are part of the DBTBD (Fig. 1A and Supp. Fig. 1-2)^18,26^. In contrast, most of PER is unstructured including its known phosphorylation domains, PER^SHORT^ and a degron (Fig. 1A)^22,23,27^. Consistent with these known structured domains, regions with consistent pLDDT scores >70 for PER were the two PAS domains, the E-helix, and two highly conserved regions of PER, namely its SYQ (613-636) and LT domains (770-794, Supp. Fig. 1-3, 1-4)^28–30^. The remaining ∼70% of PER was unstructured, consistent with previous predictions (Supp. Fig. 1-4)^31,32^.

DBT is predominantly structured with stereotypical kinase N- and C-lobes and only an unstructured C-terminus. All structured regions of DBT had similar scores, consistent with the crystal structures of CK1δ^15,33^ (Supp. Fig. 1-5).

Predicted aligned error (PAE) scores also showed high confidence in the position of the PAS domains of PER as well as the positions of the N- and C-lobes of DBT (Supp. Fig. 1-4A). We next predicted the PER/DBT complex, which identified the PER SYQ and LT domains as high confidence regions near DBT (Fig. 1B, Supp. Fig. 1-4A). Many residues within these PER domains were positioned only 2-5 Å from DBT, indicating a predicted interaction between the two proteins (Fig. 1B-C). Similar scores in these regions were calculated for human, mouse, and *Xenopus* orthologs of PER2 and CK1δ (Supp. Fig. 1-2, 1-3), indicating consistent predictions in these orthologous regions. Multiple sequence alignments between PER from *Drosophila melanogaster* and evolutionary orthologs showed that these residues and the SYQ and LT sequences have high conservation across the animal kingdom, suggesting their likely importance for PER function (Fig. 1D). The association of the SYQ and LT helices with DBT appears specific to the PER-DBT complex as there was no predicted interaction between these helices and other putative circadian protein partners (Supp. Fig. 1-4B-D). The two helices were also not predicted to interact with each other when PER was folded alone (Supp. Fig. 1-5B).

### Validating an AF-predicted Stabilizing Salt Bridge

To test the importance of putative binding residues in vivo, we altered interesting amino acids within the PER SYQ and LT helices. We began with a pair of interacting residues, E621 (SYQ) and K789 (LT); they create a stabilizing salt bridge between the two helices, which are orthogonal to each other in the predicted structure (Fig. 2A). Mutations were made in a *per*13.2 rescue plasmid, which was inserted using directed insertion into the III chromosome (see Methods). Rescue with the control plasmid using this method resulted in flies with a measured period of 22.7 hrs (Supp. Fig. 2-1), against which all mutants were compared.

**Figure 2.**
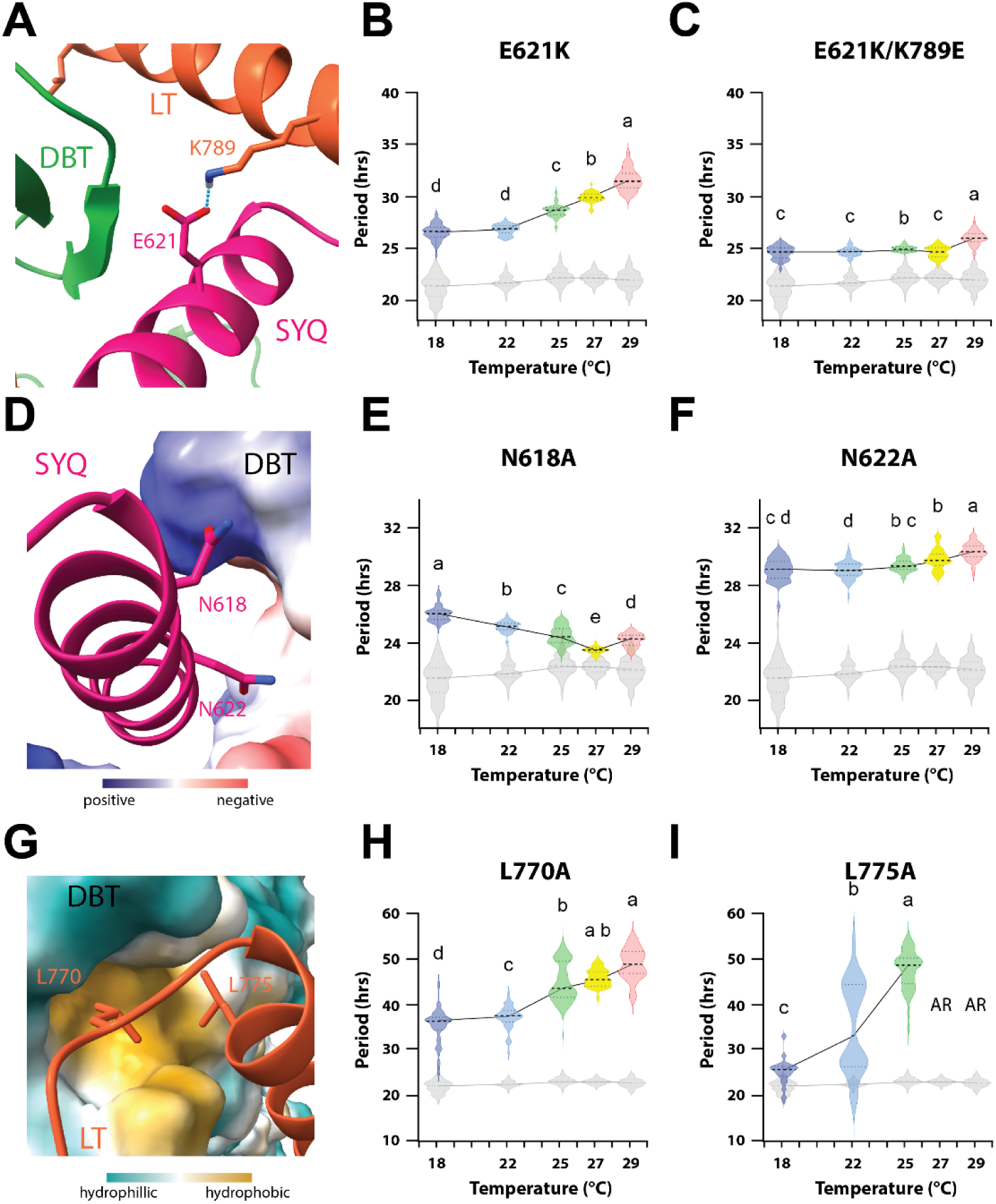
SYQ and LT mutants show significant loss of circadian temperature independence and alter the period. (A) A putative charge cluster is predicted between E621 on the SYQ domain and K789 on the LT domain. A predicted H-bond is shown as a blue dashed line between these residues. (B) Fly behavioral data for E621K shows a significant increase in period at 25 °C and a loss of temperature independence. (C) Both effects are largely rescued in the double mutant. (D) The polar residues N618 and N622 are shown to interact with charged areas adjacent to the catalytic cleft of DBT. DBT shown as surface model. (E-F) Fly behavioral data show N618A and N622A mutants have a lengthened period at 25 °C but have lost temperature independence in opposite ways. (G) The residues L770 and L775 sit in a hydrophobic pocket of DBT. DBT is shown as a surface model. (H-I) Both mutants have dramatic effects on period at 25 °C and on temperature-independence. Letters (a-d) indicate statistical differences within mutant groups (see Methods for details). WT control is shown in grey for reference.

We first decided to test the effect of charge removal and charge reversal, by creating E621A and E621K mutants respectively. Both mutants showed significant free-running period lengthening: E621K resulted in a period change of ∼5.5 hours at 25 °C (Fig. 2B), whereas E621A showed a more modest but still significant ∼45 min period increase (Supp. Fig. 2B). The difference between the two mutants is likely due to the E621K charge repulsion from the neighboring K789 residue compared to the more neutral E621A mutation (See below).

The E621K mutant also had a significant impact on the temperature independence of the circadian rhythm. We measured a period increase of 5 hours between 18 °C and 29 °C with longer periods at higher temperatures (Fig. 2B). The effect of temperature was most pronounced at higher temperatures: there was only a ∼20 min increase from 18 to 22 °C with almost all of the 5 hour increase occurring between 22 °C and 29 °C.

Strikingly, the double mutant E621K/K789E had a much-reduced period compared to the single mutant (24.8 h vs. 28.2 h @ 25 °C) and was able to restore most of the temperature independence (Fig. 2C); only a 1.4 h period increase between 27 °C and 29 °C remained. This near-complete rescue of the single mutant phenotype with a second compensating mutation supports the prediction of a stabilizing salt bridge between the E621/K789 residues and increases confidence in other aspects of the predicted structure.

### The SYQ Domain is Important for Circadian Regulation

We identified and mutated three putative SYQ interacting residues with DBT. Two asparagine mutations, N618A and N622A (Fig. 2D) significantly lengthened periods by 1.9 and 6.7 hours respectively (Fig. 2E-F). The F626A mutation resulted in arrhythmicity (Supp. Fig. 2-1D-E) and so was not included in the subsequent temperature studies described directly below.

N622A had a very modest 1.25 hour increase in period between 18 °C and 29 °C, (Fig. 2F). Interestingly, N618A had a dual effect with temperature: a 2.6 hour *decrease* in period between 18 °C and 27 °C and a smaller but still significant 45 minute increase between 27 °C and 29 °C (Fig. 2E). This suggests a different effect of temperature above 27 °C.

### Two LT Domain Mutations cause Dramatically Long and Temperature-Sensitive Periods

Deletion and mutation studies have previously illustrated the importance of the LT domain for PER binding to DBT and CK1δ, in flies and mammals respectively^24,25,34^. In addition to the K789 residue involved in the salt bridge described above, AlphaFold predicted interactions of a small hydrophobic region in the N-lobe of the kinase with two conserved LT hydrophobic residues, L770 and L775 (Fig. 2G). A previous study in mice showed that the PER2 ortholog of L775, PER2 L730, is important for kinase binding, but its effect on circadian period was complicated by the presence of unaltered PER1 and PER3 in the genetic background^34^. There are no comparable complications in flies with its singular *period* gene.

L770A and L775A both have huge effects on circadian period at the standard assay temperature of 25 °C with average free-running periods of 44.8 and 47.4 hours, respectively (Fig. 2H-I). To our knowledge, these are the longest periods from single *period* point mutants (Supp. Fig 2-2). Additionally, L770A and L775A flies had very steep temperature response curves (Fig. 2H-I): L770A had an observed change of 13.4 hours from 18 °C to 29 °C (Fig. 2H). Although L775A was arhythmic at 27 °C and 29 °C, its period changed by a remarkable 22.4 hours between 18 °C and 25 °C (Fig. 2I). L775A also has a large standard deviation in period length at 22 °C, like L770A at 25 °C, due perhaps in both cases to being close to the inflection point of the period-temperature profiles between 22 °C and 25 °C.

### Extreme Behavioral Phenotypes are due to a Lengthened Molecular Clock

To make sure that these extreme phenotypes were not behavioral oddities and to gain insight into mechanism, we assayed RNA and protein cycling with RNAseq and Western blotting from L775A fly heads. Under normal light-dark (LD) conditions, L775A flies were arhythmic and had constantly high *period* mRNA and protein (Supp. Fig. 3-1A-C), perhaps because of the dramatic difference between the entraining light cycle of 24hrs and the much longer endogenous period. In contrast, *period* mRNA undergoes cycling with about a 48 hour period in DD conditions (Supp. Fig. 3-1D), similar to its behavioral period. Because the arrhythmicity and constant high levels of PER and its mRNA in LD made it impossible to align the much longer circadian cycle of the mutant flies relative to that of wild-type flies, we adopted a modified entrainment regime to synchronize the molecular clocks: flies were exposed to 5 days of constant light (LL) to render flies arrhythmic and allow for PER degradation^35–37^, after which they were shifted into constant dark (DD) conditions (Fig. 3A). WT and L775A flies subjected to this regime showed the same DD periods as shown above and now appeared to enter DD in phase after lights-off (Fig. 3B) with DD0 ∼ ZT12 at about the beginning of transcription (Supp. Fig. 1-1).

**Figure 3.**
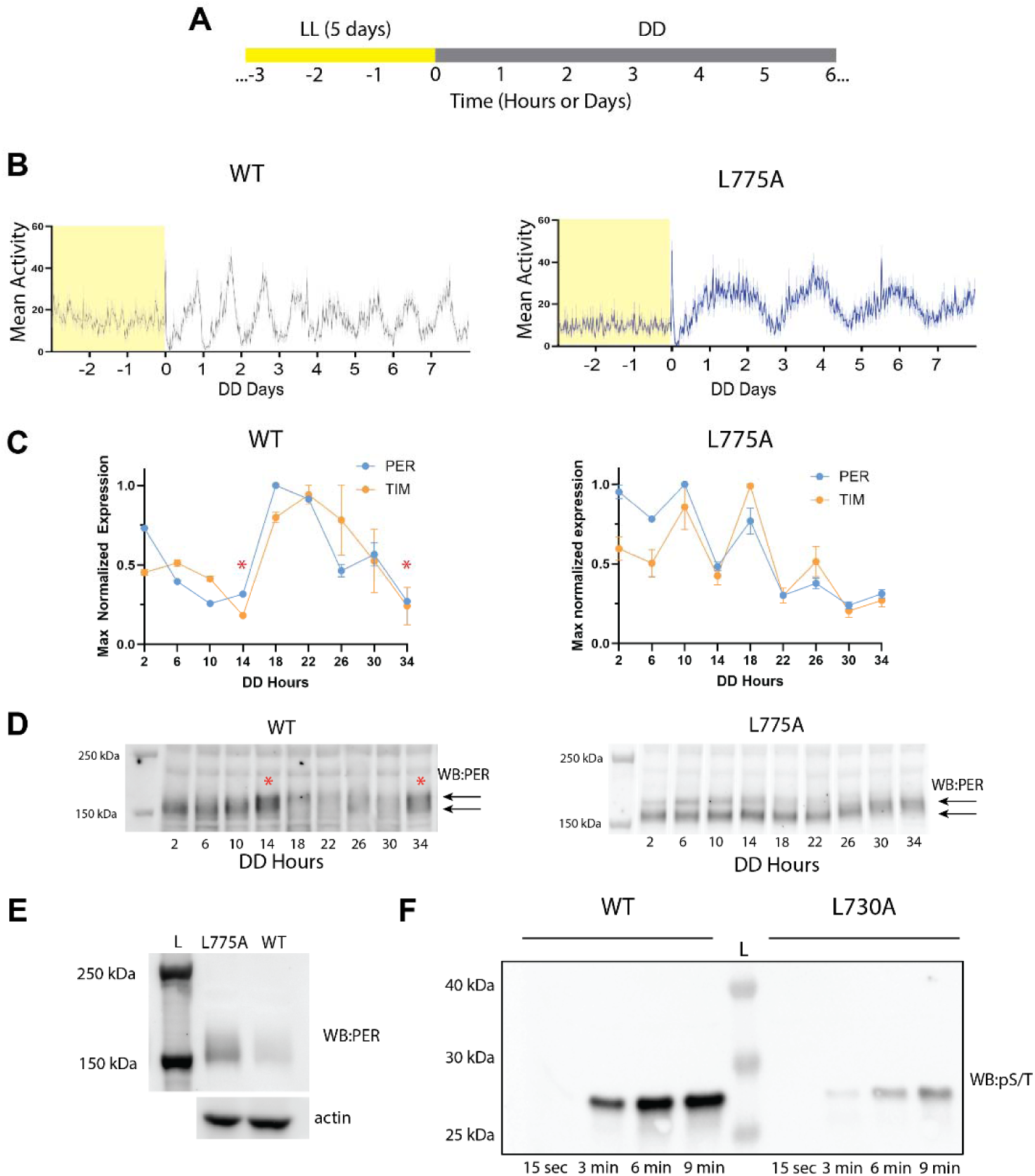
L775A mutant has slowed RNA cycling of PER and TIM (or is it fast? I see it go up and down quickly, every 4 hours??) and PER phosphorylation. (A) Diagram showing the LL to DD protocol. (B) Activity traces for WT and L775A flies after 5 days of constant light (LL) show that both genotypes are synchronized in behavior coming out of LL, and that L775A maintains its ∼48hr period of activity. (C) Max-normalized expression levels of mRNA for *per* and *tim* over 32 hours of DD. (D) Western blot from DD head timepoints show that L775A-PER levels are still high after LL and have slowed phosphorylation and degradation pattern compared to WT. Shift in apparent molecular weight in PER due to phosphorylation shown with two arrows. Red stars (C and D) show the location where high PER protein correlates with low RNA in WT as expected. (E) Western blot of DD 0 shows higher levels of PER in L775A compared to WT. (F) Western blot of anti p-Ser/Thr showing phosphorylation kinetics of 25 μM of WT or mutant hPER2 peptide by (residues: 579-751) by 1 μM CK1δ at 25 °C showing that in vitro phosphorylation of the mutant L730A-hPER by CK1δ is very slow compared to WT.

Notably, the cycling of PER-L775A RNA and protein was very different compared to WT. WT RNA and protein cycle with RNA minima corresponding to protein maxima as expected (Fig. 3C-D; red stars). In contrast, the L775A mutant showed a steady decline in RNA while protein remained relatively constant (Fig. 3C-D). Additionally, the stereotypical shift in PER migration due to hyperphosphorylation^19,38^ was greatly delayed for L775A-PER (Fig. 3D, bottom panel), suggesting that it influences the delayed RNA decline.

To further explore this putative delayed phosphorylation program of the L775A mutant, we conducted *in vitro* assays to compare the phosphorylation rates of human CKIδ:ΔC (residues 1-294) on the human CK1 binding domain of hPER2 peptide (hCK1BD; residues 579-751; including SYQ, LT and the FASP domain) vs the orthologous L775A (hCK1BD:L730A). Human CKIδ was used because of the inability to express DBT in E. coli to the required yields of soluble active protein. Kinase and peptide were combined to a final concentration of 1 μM and 25 μM respectively and the phosphorylation of the peptide was monitored at 25 °C using anti-p-S/T antibodies over nine minutes in three-minute intervals. The phosphorylation rate of the mutant hCK1BD:L730A was severely decreased compared to WT hCK1BD (Fig. 3F), revealing that the enzyme-substrate interaction via the PER CK1 binding domain helices strongly influences the rate of PER phosphorylation. This is consistent with the *in vivo Drosophila* results (Fig. 3D) and suggests that the higher L775A PER levels measured in Fig. 3E are a consequence of their lower phosphorylation rates.

### Lower Temperature Partially Rescues PER Degradation

*per* RNA and protein from heads are generally informative but come primarily from eyes and glia, which are likely not relevant to period determination. It is dictated by the ∼240 circadian neurons of which the four small lateral ventral neurons (sLNvs) are the primary pacemaker cells^39^. To assay these neurons, flies were imaged at fixed DD timepoints after lights off following 5 days of LL as done above (Fig. 3A). Levels of PER in the nucleus and cytoplasm of sLNv neurons were quantified and normalized to volume.

L775A-PER has a substantial difference both in overall protein levels and subcellular localization compared to WT-PER. Consistent with the Western blots, L775A-PER was at much higher levels compared to controls in both circadian neurons and other fly brain neurons (Fig. 3E, Supp. Fig. 4-1). Interestingly, the higher protein levels were primarily nuclear, something not observed in WT cells or expected for these timepoints (Fig. 4A-C). To determine if this was due to accelerated nuclear localization or reduced nuclear export, we imaged these cells at 30 min intervals from DD0 (lights off) to DD2 (two hours after lights off).

**Figure 4.**
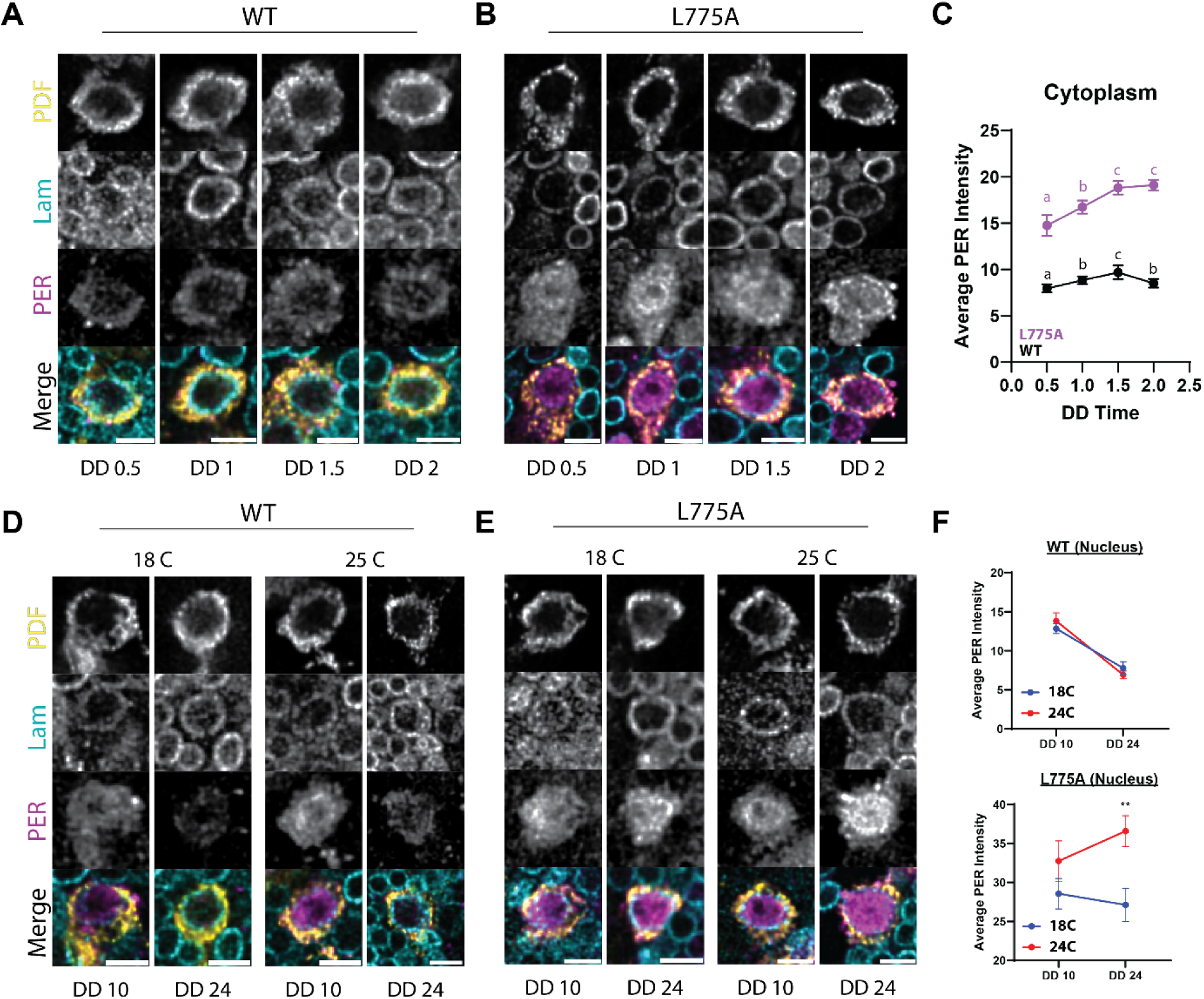
Low temperature rescues the degradation rate of PER-L775A. (A-B) During the accumulation phase both WT-PER (A) and L775A-PER (B) at 25 °C show increases in the levels of cytoplasmic protein in the sLNvs as expected for these timepoints. Anti-pigment dispersing factor (PDF) was used to label the cytoplasm of the clock cells and lamin (LAM) marks the nuclear membrane (all cells). (C) shows the quantification of the images with one-way ANOVA results shown as letters. (D, F) WT-PER displays temperature-independent PER degradation in sLNv nuclei from DD 10 to 24 as expected. In contrast, L775A-PER (E-F) reveals temperature-dependent effects as the level of L775A-PER in sLNv nuclei at 18 °C is significantly lower than that at 25 °C, quantified in (F). Stars indicate result of t-test with p-value < 0.01.

Under normal conditions, PER is cytoplasmic before accumulating substantially in the nucleus after several hours (Supp. Fig. 1-1)^40,41^. If the enhanced nuclear localization of L775A-PER was due to accelerated localization, we would have expected to see no or little accumulation of mutant protein in the cytoplasm shortly after lights-off. Yet, cytoplasmic L775A-PER increased like WT-PER over this time period despite starting at a higher overall level (Fig. 4A,C). This makes it unlikely that the high levels of nuclear L775A-PER are due to more rapid nuclear import.

Since PER degradation is also dependent on phosphorylation^42,43^, we considered that slow degradation could contribute to the enhanced protein levels in sLNv neurons as well as the long period of L775A-PER. To this end, we imaged mutant and WT-PER in these neurons at DD 10 h and then again 24 h after lights-off; this should be the PER degradation phase based on the well-known WT-PER pattern. Indeed, WT-PER levels decreased significantly during this interval (Fig. 4D, F top panel). In contrast, L775A-PER levels increased between these time points (Fig. 4E-F), a result also consistent with its much longer behavioral period.

Since the L775A mutant is temperature-sensitive and causes a much shorter period at 18 °C (Fig. 2I) we also assayed PER levels at this lower temperature. Indeed, L775A-PER levels at DD 24 were significantly lower than that at 25 °C. This pattern is closer to that of PER-WT, which had an indistinguishable pattern between 18 °C and 25 °C (Fig. 4C).

## Discussion

The DBT-binding region of *Drosophila* PER was identified and shown to be essential for rhythmicity almost 20 years ago^25^. More recent data indicates similarly that a conserved sequence domain within that region is essential for mammalian rhythmicity as well as the association between mammalian PER and CK1 in vitro^34^. These previous deletion and point-mutant studies were based solely on sequence conservation^24,25,34^, whereas our experiments here were guided by AlphaFold, which was designed to illuminate protein structure and protein-protein interactions^44,45^and has even been used to screen for novel interactors in large protein complexes^46–48^; AlphaFold generated the first high confidence atomic model of the CK1/DBT-PER interaction, some of which we successfully tested experimentally with PER mutants.

The LT domain L770A and L775A mutants resulted to our knowledge in the largest reported changes in period and temperature sensitivity measured in *Drosophila* for single amino acid substitutions; the previous record was held by the degron mutant S27A-PER (32.6 h) and TIM^rit^ (34.8 h, Supp. Fig. 2-2)^10,49^. Moreover, the data indicate that the PER SYQ domain is an additional structural element that interacts with DBT and is important for circadian period and temperature independence. Because LT and SYQ as well as many of its residues are highly conserved and AlphaFold gives similar scores for other animals (Supp. Fig. 1-3), most of these findings should be broadly applicable. In fact, we directly demonstrate that the human PER ortholog of L775A is a much less efficient in vitro substrate for CK1-mediated phosphorylation (Fig. 3F).

Many of the mutations in this study lengthen circadian period, which become even longer at higher temperatures. Longer periods are generally interpreted to reflect a slower clock due to a reduced rate at a core timing step (Supp. Fig. 1-1). The extreme period and temperature phenotypes of L770A and L775A likely reflect the importance of these hydrophobic residues for the association between CK1/DBT and PER. Taken together with other long period mutants in the DBTBD, which become even longer at higher temperature (Fig. 2), they indicate two general principles: 1) The conserved interface^18^ between the kinase and the PER SYQ and LT domain helices enhances the rate of phosphorylation (Fig. 2-4); 2) Higher temperatures further weakens the CK1/DBT-PER association (Fig. 2B, H-I). Perhaps the complete loss of rhythmicity of the L775A mutant at high temperatures reflects a complete or near-complete loss of DBT binding like the previous LT-KO studies^25^.

These principles suggest that the temperature insensitivity of kinase-mediated PER phosphorylation may be the result of two features at higher temperatures: an enhanced dissociation rate between the kinase and PER, which counteracts an increased catalytic rate. This possibility complements previous mechanisms of temperature independence, which are principally based on the timing of direct phosphorylation events and have often assayed smaller effects than those observed here ^14,15,17,50,51^. Future biophysical experiments should be able to confirm the importance and magnitude of the temperature effects on the dissociation rates of the kinase with wild-type and mutant substrates.

L775A-PER also had significant and unexpected localization defects. There are previous reports of DBT-mediated phosphorylation effects on PER localization albeit with somewhat conflicting results^25,52^. Localization has also been shown to contribute to circadian temperature-independence^53^, although the underlying mechanism is unclear. Nonetheless, the more poorly phosphorylated L775A mutant clearly has enhanced nuclear levels, which is consistent with previous work using the catalytically inhibited DBT^AR^ mutant^52^. A cause of slow phosphorylation (Fig. 4) is also consistent with recent mammalian CK1δ cell culture experiments^54^. While our study has helped solidify the importance of the kinase-PER interaction for reducing nuclear levels, future studies will be needed to clarify the mechanisms that regulate PER export as well as its degradation.

Although the LT domain mutant effects can be mostly easily interpreted as reducing the interaction with DBT, the understudied SYQ domain has more complicated effects. SYQ was first recognized as a short sequence in the *C. elegans* heterochronic gene lin-42, which is highly conserved with eukaryotic PER^30^. SYQ has also been previously implicated in kinase binding and phosphorylation through large deletions and chimeric proteins in mammalian systems^28,29^, but these experiments included the LT domain making the individual contribution of SYQ uncertain. Although our new data exploring point mutations in the SYQ domain indicate that this helix is a substantial contributor to DBTBD function in the regulation of circadian period and temperature-independence, the independence of SYQ from LT – and vice versa – has not yet been assayed.

SYQ mutations may also cause more complicated phenotypes than those in LT. The simplest SYQ mutant is F626A (Supp. Fig. 2-1E); its arrhythmicity suggests that loss of this hydrophobic residue severely destabilizes the DBTBD-DBT interaction. The N622A mutant has a long period but does not change much with temperature (Fig. 2F). In contrast, the N618A mutant has a much more modest period effect, but it becomes both shorter and longer with increasing temperature (Fig. 2E). Given the predicted position of the SYQ-domain helix across the junction between the kinase N- and C-lobes, these phenotypes may emerge from a mix of effects: one might be on the stability of the PER-kinase interaction and the other from allosteric effects on kinase activity. In any case, it is certain that future biochemical, biophysical and structural studies will be essential to determine how SYQ as well as LT interact with DBT/CK1 and contribute to the regulation of circadian period and its temperature-independence.

## Acknowledgements

We would like to thank Andrew Stone and the Brandeis Light Microscopy Facility (RRID:SCR_025892) for the use of SVI Huygens and Imaris to process and quantify our confocal data. This study was supported by funding from the Howard Hughes Medical Institute.

## Materials and Methods

### AlphaFold

Five different seeds were used to predict five structures per seed for a total of 25 predicted PER-DBT dimers using AlphaFold 2.3 multimer run on a local server. All structures were run without templates. These structures were then analyzed for regions with consistently high predicted local distance difference test (pLDDT) scores and low predicted aligned error (PAE) scores to identify regions most likely to be biologically accurate.

Alignments were performed by Clustal Omega between PER sequences from *Drosophila melanogaster* (P07663), *Homo sapiens* (O15055, O15534), *Mus musculus* (O54943, O70361), *Xenopus tropicalis* (B2GUA4), *Caenorhabditis elegans* (Q65ZG8), *Apis cerana* (Q2PHK2), *Branchiostoma floridae* (A0A9J7M8T4), *Panulirus ornatus* (UPI003A85C58F), *Eurytemora carolleeae* (UPI000C774DD3), and *Callorhinchus milii* (UPI001C3F6682). Consensus sequence values were generated in ChimeraX 1.10 in MultAlign Viewer using the AL2CO algorithm using default parameters.

### Fly Stocks and Rearing

Flies were raised on standard cornmeal medium supplemented with yeast at 25 °C under 12h light: 12h dark conditions.

### Generation of Fly Lines

Mutations were made in pAttB-Per(13.2)-HA-HIS rescue plasmid (gift from Joanna Chiu, UC Davis). Sequencing-verified plasmids were injected into the third-chromosome VK00027 site by BestGene, Inc (Chino Hills, CA, USA). Successful transformants were screened by eye color.

Mutant flies were then crossed to virgin Per01 flies and F1 males were collected and assayed for behavior. Using this paradigm, WT control flies showed a slightly decreased period at 25 °C, likely due to insertion site effects (see Supp. Fig. 2-1A). This breeding scheme allows for the stoichiometry of PER to be held constant (one active allele/fly) since increasing the copy number is known to have circadian behavioral effects^55,56^

### Circadian Behavior Assay

For behavior analysis of flies, 0-5 day old males were collected and aged until ∼ 2 weeks old using rearing method above with food changes every 3-4 days. Male flies were loaded into Drosophila Activity Monitor (DAM) tubes under standard conditions, i.e., sucrose food (4% sucrose and 2% agar). Flies were entrained to at least 3 days of a 12 hour light: 12 hour dark cycle (LD) before switching to constant darkness for at least 12 days. Circadian free-running periods were determined using the CWT analysis in the Rethomics package in R^57^.

For LL to DD experiments, flies were entrained for 3 days in LD at the indicated temperature before being switched to LL at the end of the light period of day 4. Flies were subjected to 5 days of LL, after which they were switched into DD. Flies were collected at the DD timepoints noted.

### Generation of Sequencing Libraries

For RNA cycling, L775A mutant and WT flies were entrained at 25 °C for 5 days and then frozen at 4 hr. intervals over one day of LD and the first two days of darkness (DD 1-2). RNA from fly heads (10-15) was extracted using TRIzol reagent (ThermoFisher #15596026). Libraries were generated using the xGen RNA Library Prep Kit (IDT #10009814) using mRNA isolated using the NEBNext Poly(A) mRNA Isolation Kit (NEB #E7490). Libraries were purified using AMPure XP Beads (Beckman Coulter #A63881) and quantified using a High Sensitivity D5000 ScreenTape (Agilent # 5067-5592) on a TapeStation.

These libraries were then sequenced on Illumina NovaSeq6000 at Novogene Corporation Inc. (Sacramento, CA, USA) generating 20-38million total raw paired end reads (2*150bp) per sample.

### Sequencing Data Analysis

Pair-end reads were pseudoaligned and quantified using Salmon^58^. Count tables were imported into R using Tximeta^59^ and transcript-level counts were summarized to gene-level counts. Gene-level counts were normalized using DESeq2^60^.

### qPCR

Fly heads were collected on dry ice and manually lysed using pestle in Trizol (Invitrogen #15596026) followed by flow through a Qiashredder (Qiagen # 79656). Chloroform was added and RNA was separated using a Phasemaker tube (Invitrogen #A33248). RNA was precipitated using isopropanol with Glycoblue (Invitrogen #AM9515) by placing at -20 °C for at least 1 hour and pelleting for 30min at 21,000 g. The pellet was cleaned with 80% ethanol 2X after which it was dried and resuspended in IDTE. DNA was removed using the TURBO DNA-free system (Invitrogen #AM1907), and the RNA concentration was measured.

cDNA was created using 400 ng of RNA with 100 U of Maxima H-minus RT (Thermo Scientific # EP0752) using OligodT primers under standard conditions.

qPCR was performed in triplicate using SyGreen PCR master mix (PCR Biosystems #PB20.14-01) using 1ul of 10 nmol forward and reverse primers. cDNA was diluted 1:10 and 1ul was used for the final reaction. Primers used were published previously^61^.

### Confocal Imaging

For confocal imaging, flies were collected at the times indicated and were fixed with PFA at 4C for ∼ 6 hours after which brains were dissected. They were blocked at room temperature using 5% NGS in PBS supplemented with 0.5% Triton-X 100 (S-PBST) for 3 hours, after which the brains were incubated in primary antibody mixture (antibody + S-PBST) for 2 days at 4 °C . The brains were washed 3 times with PBST (30 min each) and then incubated with secondary antibody mixture for 3 days at 4 °C. They were then washed as above, and then mounted onto slides separated into groups using Secure-Seal spacers (Invitrogen #S24737) and SlowFade Glass (Invitrogen #S36917) and covered using #1.5 coverslips. Primary antibodies: mouse IgG2b anti-PDF (DSHB #PDF-C7; 1:1000); mouse IgG1 anti-Lamin (DSHB #ADL101; 1:1000); rabbit anti-PER (custom antibody, 1:500). Secondary antibodies: goat anti-mouse IgG1 AF488 (Invitrogen #A21121); goat anti-mouse IgG2b AF568 (Invitrogen #A21144); goat anti-rabbit AF647 (Invitrogen #A21245).

Imaging was performed on a Leica Stellaris 8 confocal microscope using a white light laser (85%) using a 63x/1.4 oil objective with the following three channels: 1)Excitation of 499nm at 7% detected using a HyD S collecting in counting mode from 488-568nm with TauGating from 0.8 to 2.4ns; 2) Excitation of 579 at 0.5% detected using a HyD S collecting in analog mode from 584-651nm with a Gain of 2.5; 3) Excitation of 653nm at 2.8% intensity using a HyD X detector collecting in counting mode from 663-750nm with TauGating from 1.4-4.1ns. Pixel dwell time was 1 μs with a voxel size of 86×86×300nm.

### Image analysis

Images were processed using SVI Huygens deconvolution software to improve signal-to-noise ratio before final quantification using Imaris (v. 10.2 for Core Facilities). For quantification, surfaces were created in 3D using PDF and lamin channels to identify the soma and nucleus of the cells respectively. These were then imported into the Imaris Cells module for quantification of PER. All analysis was done in 3D with average voxel intensities used for quantification unless otherwise noted.

### Protein production

#### hCK1δ:ΔC

His6-GST-SUMO-hCK1δ:ΔC (kinase from Homo sapiens; UniProt: P48730•KC1D_HUMAN; residues 1-294) was codon-optimized for E. coli K12 and cloned in pET28(+) (NdeI & XhoI sites, performed by TWIST Biosciences). E. coli BL21 (DE3) cells (NEB) were transformed with said plasmid according to manufacturer’s instructions. A starter culture (50 mL of LB media, 50 µg/mL kanamycin) was inoculated with a single colony and incubated overnight at 37 °C and 220 rpm (Multitron Standard, INFORS HT). To set up expression cultures, 1 L of LB media (50 µg/mL kanamycin) was inoculated with 5 mL of starter culture. Cells were cultured at 37 °C and 220 rpm until they reached an OD*_600_* of 0.6 – 0.8 and were induced with 0.1 mM IPTG. The temperature was subsequently lowered to 23 °C and the expression cultures were incubated overnight (∼18 h). Cell pellets were harvested via centrifugation at 4,000 rpm (Avanti JXN-26 & JLA-8.1000; both Beckman Coulter) for 15 min at 4 °C and subsequently stored at -20 °C or resuspended at 4 °C in buffer A (500 mM NaCl, 50 mM TRIS, 1 mM TCEP, pH 7.5) + 1 mg/mL HEW lysozyme (Sigma) and 5 µg/mL DNase (Sigma). The resuspended cells were sonicated on ice in a 4 °C cold room using Qsonica sonicator (15 s on 45 s off, total 12 min on, 45% amplitude, total time 48 min) and then clarified via centrifugation for 30 min at 18,000 rpm (Avanti JXN-26 & JA-25.50; both Beckman Coulter) at 4 °C. Supernatant was decanted and filtered (0.22 µm *syringe filter, Cytiva*) then loaded (1 mL/min Äkta Pure (Cytiva)) onto a 5 mL GSTrap™ HP (*Cytiva*) equilibrated in buffer A. CK1δ:ΔC was eluted with buffer B (500 mM NaCl, 50 mM TRIS, 10 mM glutathione, 1 mM TCEP, pH 7.5). The elution was buffer-exchanged into buffer A using a HiPrep 26/10 desalting column (Cytiva) and then digested with Ulp-1 at 4 °C overnight. After digestion, CK1δ:ΔC was loaded onto a tandem 5 mL HisTrap HP column (Cytiva) and 5 mL GSTrap™ HP column (Cytiva). Cleaved CK1δ:ΔC was eluted with 50 mM Tris, 500 mM NaCl, 50 mM imidazole, 1 mM TCEP, pH 7.5. The elution was pooled and buffer-exchanged into buffer A using a HiPrep 26/10 desalting column (Cytiva). CK1δ:ΔC was then aliquoted, flash frozen in liquid N*_2_* and stored at -80 °C. During purification all samples and buffers were kept on ice or at 4 °C (during centrifugation, sonication and FPLC application)

#### hCK1:BD (WT) & hCK1:BD:L730A

His6-NusA-TEVsite-hCK1BD (PER2 from Homo sapiens; UniProt: O15055•PER2_HUMAN; residues 579-751; WT and L730A) was codon optimized for E. coli K12 and cloned in pET28(+) (NdeI & XhoI sites, performed by TWIST Biosciences). E. coli BL21 (DE3) cells (NEB) were transformed with said plasmid according to manufacturer’s instructions. A starter (50 mL of LB media, 50 µg/mL kanamycin) was inoculated with a single colony and incubated overnight at 37 °C and 220 rpm (Multitron Standard, INFORS HT). To set up expression cultures, 1 L of LB media (50 µg/mL kanamycin) was inoculated with 5 mL of starter culture. Cells were cultured at 37 °C and 220 rpm until they reached an OD600 of 0.6 – 0.8 and were induced with 0.1 mM IPTG. The temperature was subsequently lowered to 23 °C and the expression cultures were incubated overnight (∼18 h). Cell pellets were harvested via centrifugation at 4,000 rpm (Avanti JXN-26 & JLA-8.1000; both Beckman Coulter) for 15 min at 4 °C and subsequently stored at -20 °C or resuspended at 4 °C in buffer A (500 mM NaCl, 50 mM TRIS, 1 mM TCEP, pH 7.5) + 1 mg/mL HEW lysozyme (Sigma) and 5 µg/mL DNase (Sigma). The resuspended cells were sonicated on ice in a 4 °C cold room using Qsonica sonicator (15 s on 45 s off, total 12 min on, 45% amplitude, total time 48 min) and then clarified via centrifugation for 30 min at 18,000 rpm (Avanti JXN-26 & JA-25.50; both Beckman Coulter) at 4 °C. Supernatant was decanted and filtered (0.22 µm syringe filter, Cytiva) then loaded (1 mL/min Äkta Pure (Cytiva)) onto a 2x 5 mL HisTrap HP (Cytiva) equilibrated in buffer A. hCK1BD fusion protein was eluted with buffer B (500 mM NaCl, 50 mM TRIS, 250 mM imidazole, 1 mM TCEP, pH 7.5). The elution was buffer-exchanged into buffer A using HiPrep 26/10 desalting column (Cytiva) and then digested with SuperTEV protease^62^ at 4 °C overnight. After digestion, the sample was loaded onto 2x 5 mL HisTrap HP column (Cytiva) equilibrated with buffer A. Cleaved hCK1BD was collected with the flow-through. The flow through was concentrated and subjected to size-exclusion chromatography using an S200 16/600 pg column (Cytiva) at 0.5 mL/min equilibrated with buffer A. Pooled fractions were then concentrated, aliquoted, flash frozen in liquid N2 and stored at -80 °C. During purification all samples and buffers were kept on ice or at 4 °C (during centrifugation, sonication and FPLC application).

### In vitro discontinuous kinase assay

Reaction kinetics were observed via phospho-Ser/Thr western blot. Phosphorylation of 25 µM hCK1BD or hCK1BD:L730A (hPER2 residues: 579-751; containing spanning SYQ-motif, FASP region, and LT-motif) at 25 °C was catalyzed by 1 µM hCK1δ:ΔC and performed in reaction buffer (500 mM NaCl, 50 mM TRIS, 1 mM TCEP, pH 7.5), with 0.6 mg/mL BSA (Sigma) and 5 mM Mg2+•ATP (Sigma). A reaction mixture lacking 5 mM Mg2+•ATP was incubated for 10 min at 25 °C. Mg2+•ATP was used to initiate the reaction. Samples were taken at time points: 30 s, 3, 6 and 9 min by quenching reaction aliquots with 1:1 reducing NuPage 4x LDS sample buffer (Invitrogen). Samples were incubated for 5 min at 95 °C and loaded on an SDS-PAGE (2 µL per lane) on a gel (Bolt 4-12% Bis-Tris Plus WedgeWell, Invitrogen) and run according to manufacturer’s instructions. The gel was briefly washed in MQ water and then dry-transferred (7 min) using an iBlot2 device with iBlot 2 NC Regular Stacks (both Invitrogen). Membrane was briefly rinsed in MQ water and blocked at room temperature in “Blocking Buffer for Fluorescent Western Blotting” (Rockland). Primary antibody: 1:5,000 dilution anti-pS/T polyclonal Ab (rabbit, PhosphSolutions) incubated overnight at 4 °C in “SuperSingal Western Blot Enhancer” (Thermo Scientific). Membrane was washed 3x 5 min in 1x TBS-T buffer. 1:10,000 dilution of secondary (fluorescent) antibody: IRDye800CW goat anti-rabbit IgG (LI-COR) in Intercept Ab diluent (LI-COR) was used to incubate the membrane for 1 h at room temperature. The membrane was imaged using a ChemiDoc (Bio-Rad) and DyLight 800 (4×4) setting (exposure time: 69 s).

### Statistical Analysis

All behavior was analyzed using the Brown-Forsythe and Welch one-way ANOVA with Dunnett’s T3 multiple comparisons test in GraphPad Prism. Comparative one-way ANOVAs for imaging were performed with a Tukey test for multiple comparisons.

