## Supplemental Figures for "The dramatic impact of the PER-DBT interaction on circadian timekeeping and temperature compensation"

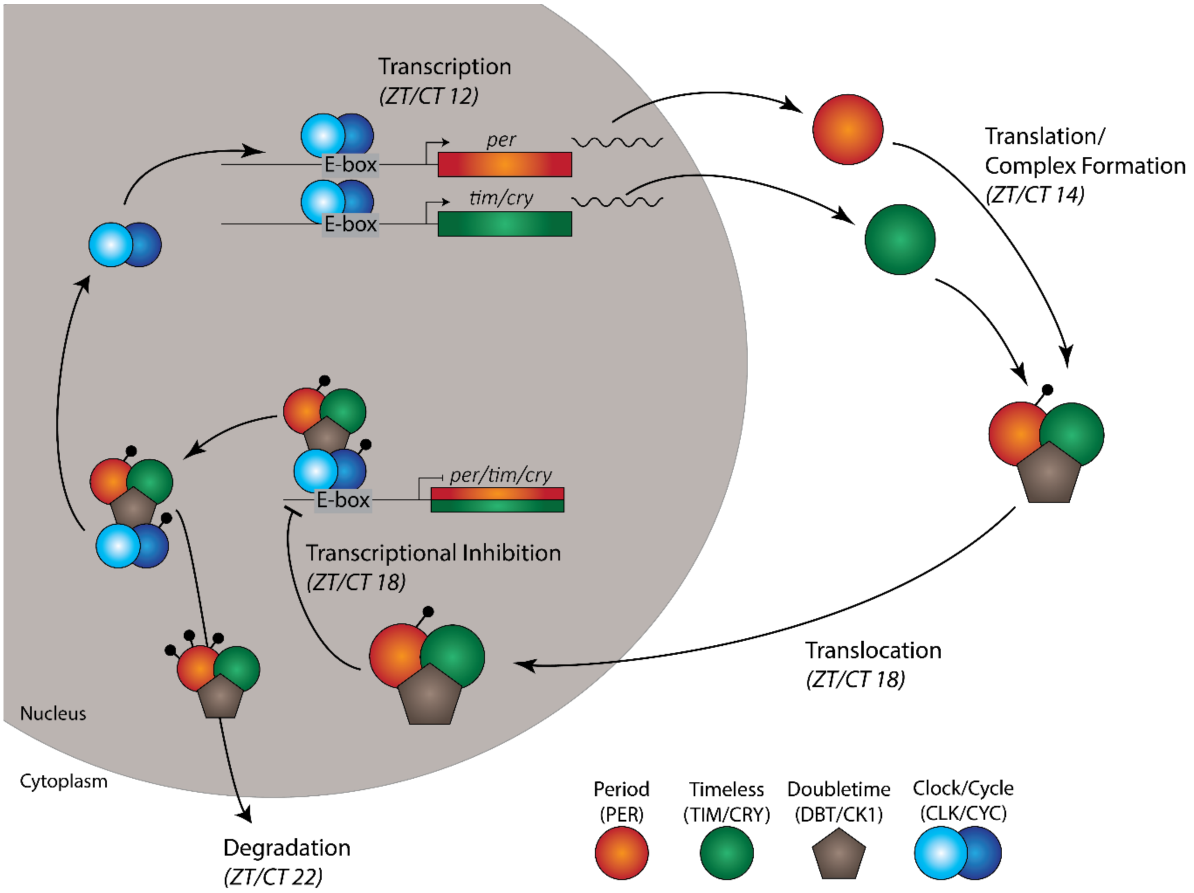


**Supp. Figure 1-1. The canonical circadian transcription/translation feedback loop (TTFL).** A graphical representation of the TTFL. Proteins are identified with both common *Drosophila* and mammalian designations for PER, TIM/CRY, and DBT/CK1. Approximate start times for each step of the clock are given underneath each major step (ZT – zeitgeber time; CT – circadian time) with ZT 0 representing lights on and ZT 12 representing lights off. CT time is time in constant darkness and is when circadian periods are measured. In wildtype flies, each step happens at roughly the same ZT or CT time each 24-hour period.

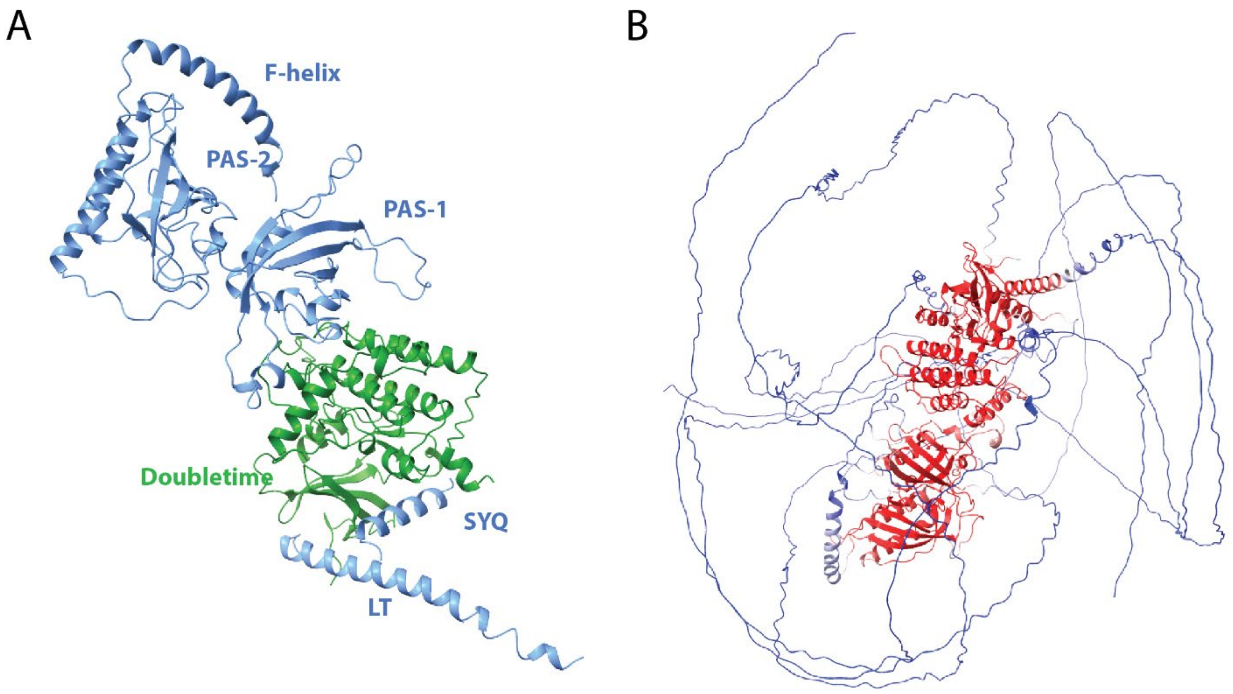


**Supp. Figure 1-2. Globular structures and pLDDT scores of full-length *Drosophila* PER/DBT AlphaFold prediction.** (A) AlphaFold prediction of the PER/DBT complex with the structured regions of DBT (green) and Period (blue) shown only; major domains are labeled. (B) pLDDT scores mapped onto a representative PER/DBT predicted complex show high levels of confidence for the globular domains of PER and DBT (red) but not for the unstructured regions (blue).

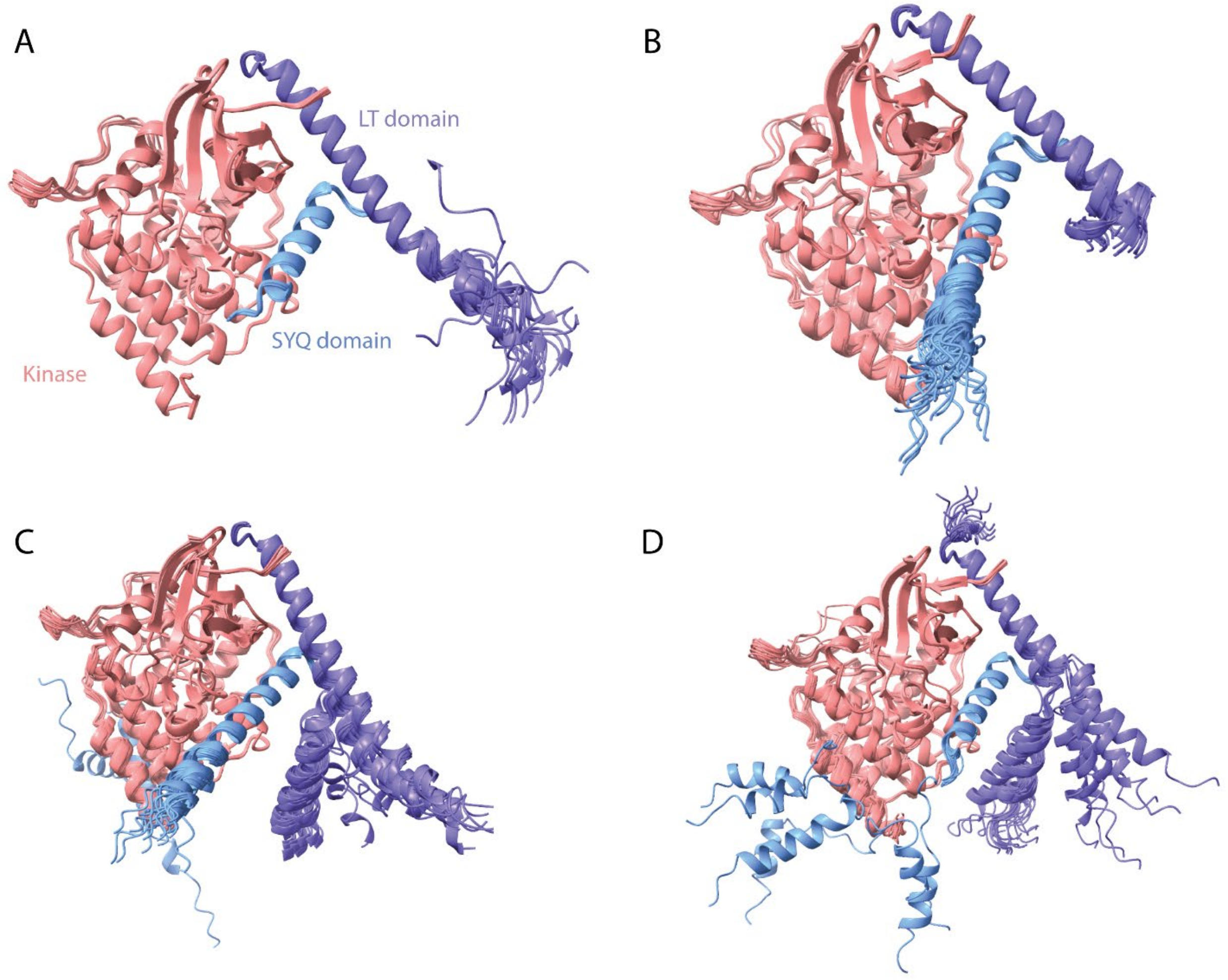


**Supp. Figure 1-3. Ensemble alignments for Period LT and SYQ domains/Kinase AlphaFold predictions.** 25 AlphaFold predicted structures were aligned for *Drosophila* (A), human (B), mouse (C), and *Xenopus* (D) using the species-specific sequence for DBT or CK1δ with their respective PER or PER2 sequences. Alignments were performed using the kinase as a reference for all the structures. Across species the LT (purple) and SYQ (blue) helices show consistent placement relative to the kinase. Only the mouse (with 1) and the *Xenopus* (with 6) have some predicted binding locations significantly different than the rest of the ensembles.

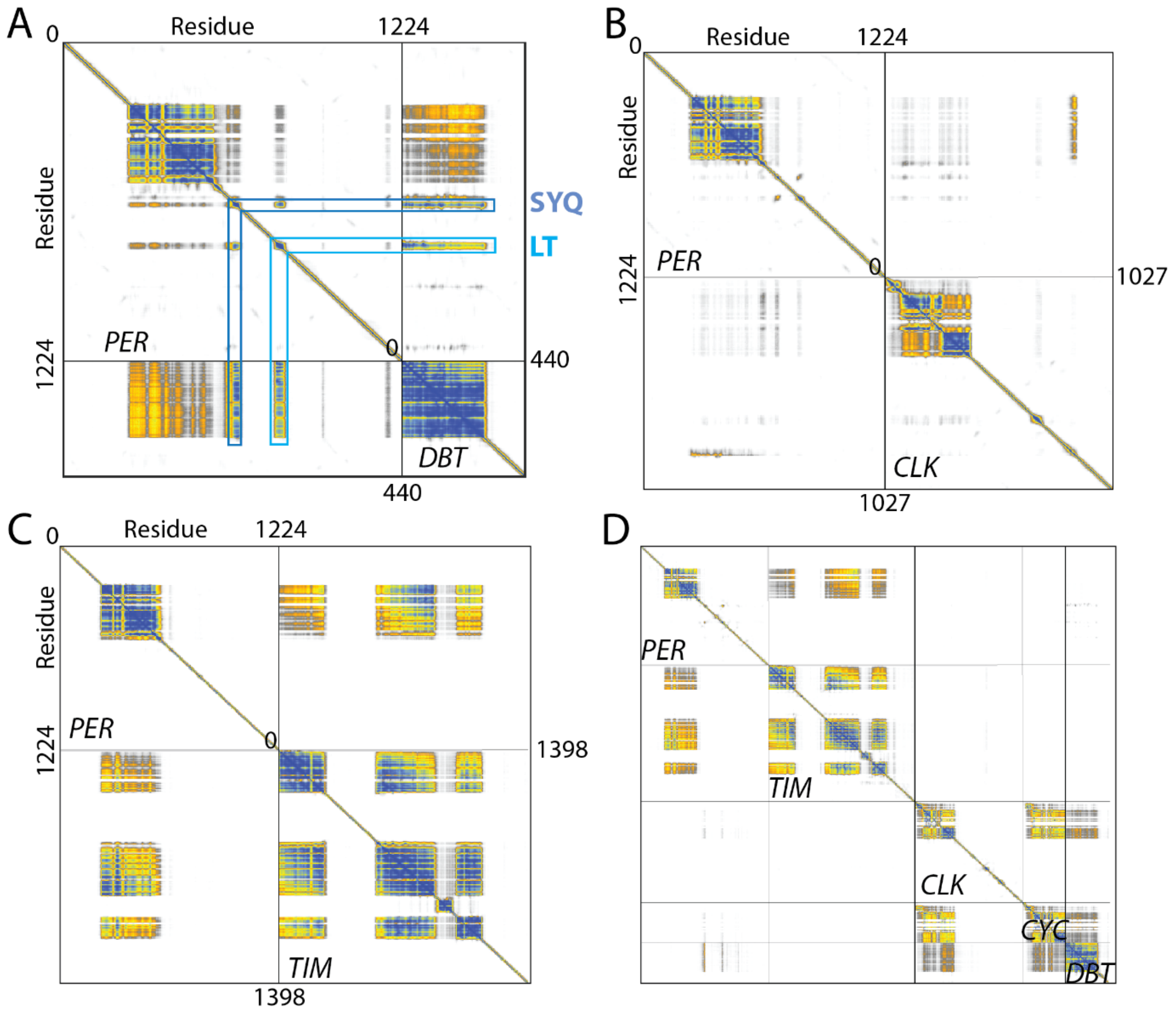


**Supp. Figure 1-4. Pairwise PAE plots of PER with circadian proteins.** Representative PAE plots of PER with circadian binding partners DBT (A), CLK (B), TIM (C), and as a whole complex (D). In (A) the location of the SYQ and LT domains are highlighted with blue bars to show the areas of high PAE scores. In (D) residue numbers for each protein match those in other panels. CYC residues go from 0-413.

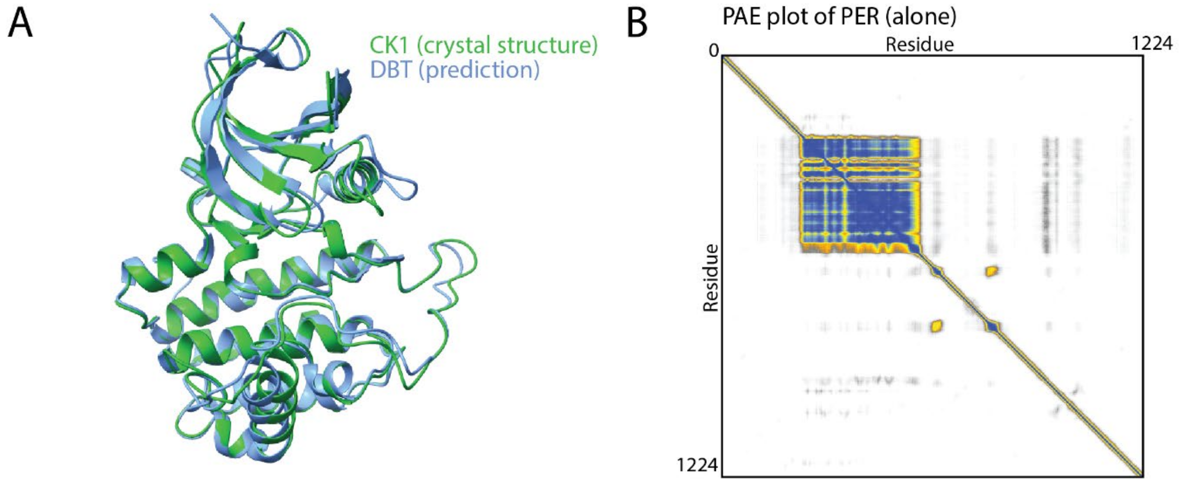


**Supp. Figure 1-5.** (A) Predicted structure of DBT (blue) overlaid with the known structure of CK1 (green, PDB 6PXO). (B) Representative PAE plot of PER alone showing that the LT and SYQ domains have only a low-confidence prediction of interaction across seeds.

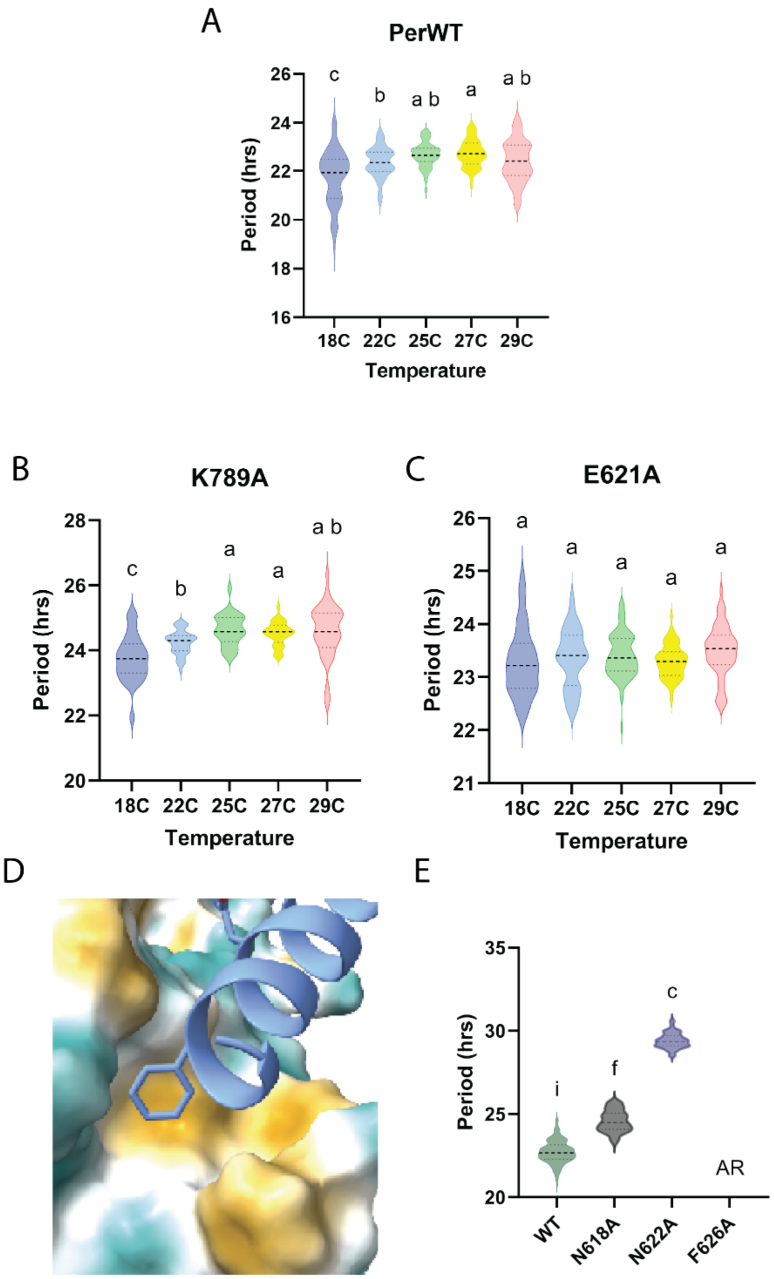


**Supp. Figure 2-1. Free-running periods for K789A and E621A from 18C to 29C.** (A) Free running period of WT flies from 18 to 29 ºC (B) The effect of temperature on the K789A mutant was small but significant. (C) The E621A mutant had no significant effect from temperature. (D) Location of F626A mutation on PER (blue) against a hydrophobic surface model of DBT. (E) Free running periods of mutants as assayed at 25 ºC showing F626 is arhythmic. Small letters indicate statistical differences within mutant groups (see Methods for details).

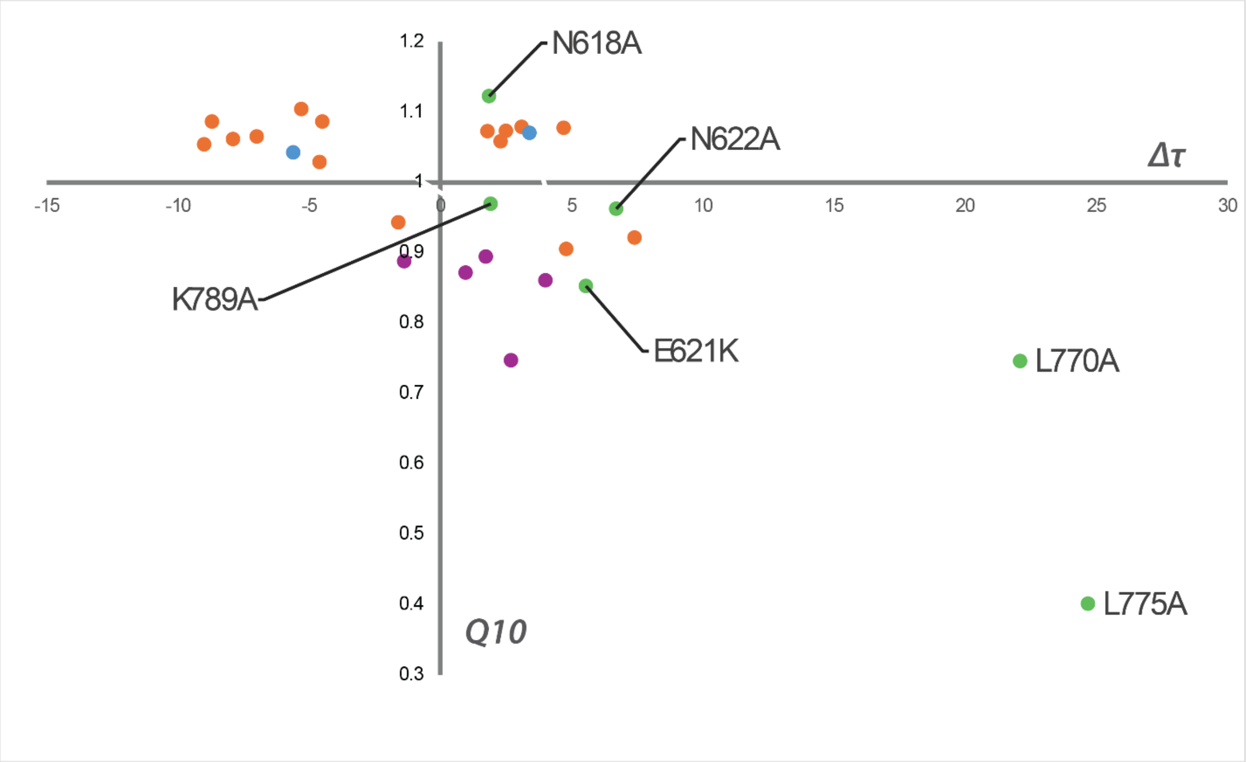


**Supp. Fig 2-2. Summary of Drosophila temperature dependent mutants.** A graph of change in free-running period (x-axis) vs calculated Q10 value (y-axis). Previous PER mutants are in orange, DBT mutants in blue, and TIM mutants in purple. The PER mutants in this paper are green and are labeled.

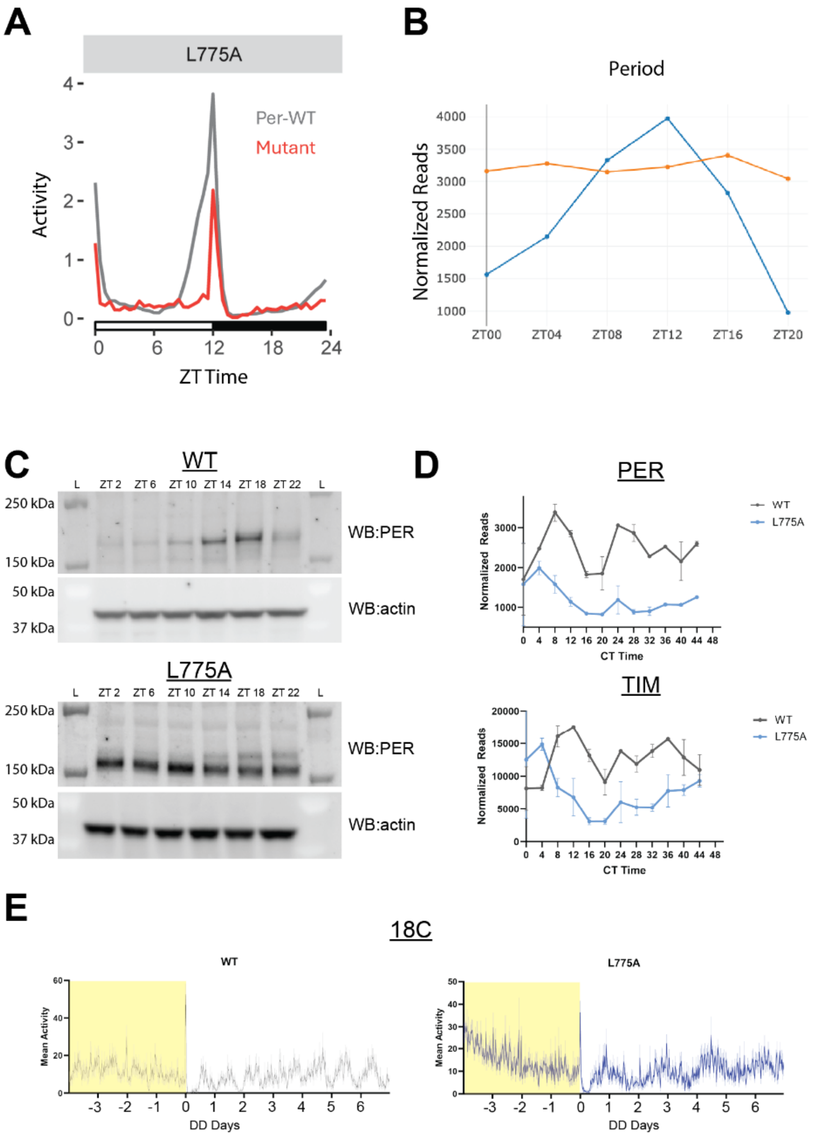


**Supp. Fig. 3-1. LD and DD data for L775A mutant**. (A) L775A flies are arrhythmic under LD conditions. (B) RNA levels in LD remain high in L775A mutant (orange) compared to WT (grey). (C) Western blots from heads show much higher levels of PER-L775A around the clock in LD. (D) RNAseq from heads (CT 0-48) show L775A-PER cycles with a period comparable to the behavior data. (E) LL to DD behavior at 18C shows a slightly shorter period in L775A as expected from behavior.

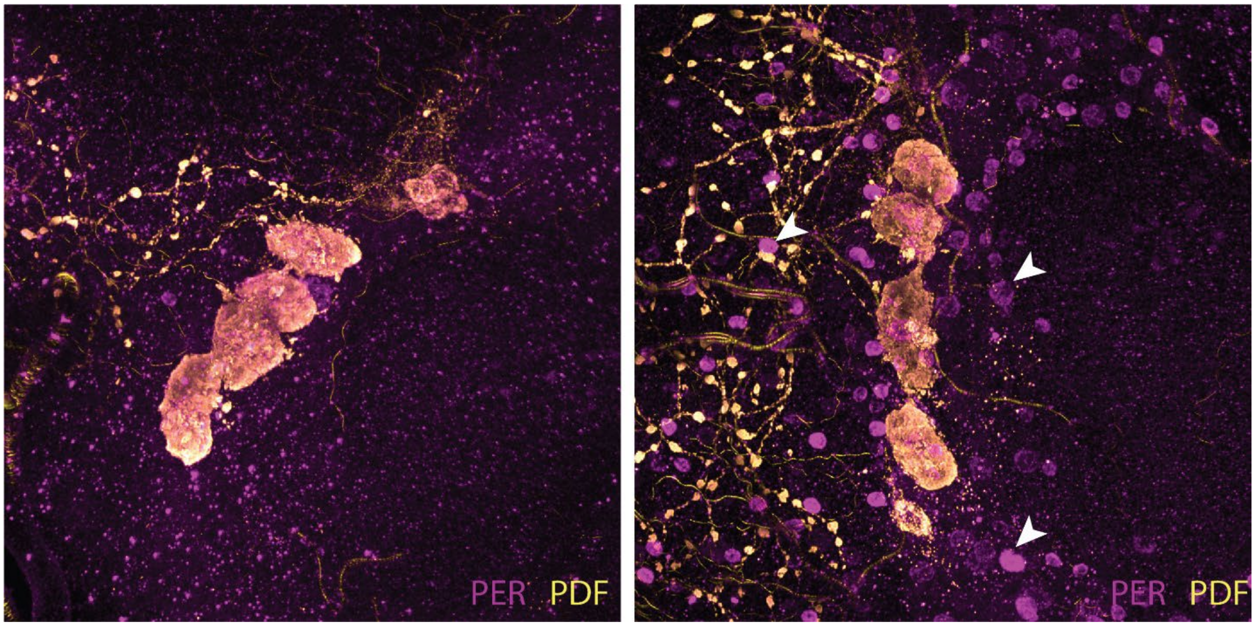


**Supp. Fig 4-1. L775A mutant results in high nuclear PER protein in circadian and ectopic cells**. Representative images of brains at DD2. PDF labeling identifies the LNv cell bodies and neurites. L775A brains (right) show many more ectopic nuclei (examples shown using white arrows) than WT (left). Image intensities were auto thresholded for each channel to best show signal.
